# Step selection analysis with non-linear and random effects in mgcv

**DOI:** 10.1101/2024.01.05.574363

**Authors:** Natasha Klappstein, Théo Michelot, John Fieberg, Eric Pedersen, Chris Field, Joanna Mills Flemming

## Abstract

Step selection analysis is used to jointly describe animal movement patterns and habitat preferences. Recent work has extended this framework to model inter-individual differences, account for unexplained structure in animals’ space use, and capture temporally-varying patterns of movement and habitat selection.In this paper, we formulate step selection functions with penalised smooths (similar to generalised additive models) to unify new and existing extensions, and conveniently implement the models in the popular, open-source `mgcv` R package. We explore non-linear patterns of movement and habitat selection, and use the equivalence between penalised smoothing splines and random effects to implement individual-level and spatial random effects. This framework can also be used to fit varying-coefficient models to account for temporally or spatially-heterogeneous patterns of selection (e.g., resulting from behavioural variation), or any other non-linear interactions between drivers of the animal’s movement decisions. We provide the necessary technical details to understand several key special cases of smooths and their implementation in `mgcv`, showcase the ecological relevance using two illustrative examples, and provide R code (available at https://github.com/NJKlappstein/smoothSSF) to facilitate the adoption of these methods. This paper is a broad overview of how smooth effects can be applied to increase the flexibility and biological realism of step selection analysis.

## 1 Introduction

Step selection functions (SSFs) are increasingly used to describe animal movement and habitat selection (Rhodes et al., 2005; Forester et al., 2009; Avgar et al., 2016). An SSF measures how an animal selects habitat at the scale of the observed movement step, whilst simultaneously estimating distributions of step lengths and turning angles. Compared to landscape-level habitat selection models (e.g., resource selection functions; RSFs), the temporal structure of SSFs better accounts for autocorrelation in animal tracking data. SSFs also make it possible to assess time-varying patterns of selection (Forester et al., 2009; Richter et al., 2020) as well as movement-habitat interactions that may be associated with behavioural patterns or landscape resistivity (Avgar et al., 2016). Their dynamic formulation makes them an important tool in predicting animal space use, with methods recently developed to scale SSF results to large-scale distributions (through simulations and mathematical methods; Signer et al., 2017; Potts and Börger, 2022; Signer et al., 2023). Therefore, to best understand animal spatial ecology, many methodological extensions of SSFs have been proposed to improve their biological realism and flexibility.

One important focus of SSF research has been to relax the common assumption that the parameters driving movement and habitat selection are constant through time, to account for behavioural variation. This can for example be done with hidden Markov models, where animals switch between behavioural states, each characterised by different patterns of movement and habitat preferences (Nicosia et al., 2017; Klappstein et al., 2023; Pohle et al., 2023). Another approach is to include interaction effects within an SSF, to specify covariate-dependent selection parameters. For example, habitat selection can vary with time (Forester et al., 2009; Richter et al., 2020), and movement speed can change with environmental resistivity (Avgar et al., 2016). Although this approach does not explicitly model behavioural switching, it provides a very flexible framework to capture time-varying movement and selection patterns. This method could be further extended to allow for non-linear habitat preferences and non-linear interaction effects (e.g., seasonal selection patterns). These ideas have been explored for RSFs (e.g., McCabe et al., 2021; Dejeante et al., 2023), but not yet for SSFs, which have the ability to assess variability in both habitat selection and movement.

Another direction of recent research has been to include random effects in SSFs, to capture variability otherwise unexplained by the model. Random slopes have been explored to account for inter-individual variability in movement and habitat selection patterns, attributable to animal “personality” and affected by phenotypic plasticity (Duchesne et al., 2010; Muff et al., 2020; Chatterjee et al., 2023). Similarly, spatial random effects can help account for spatial pattern or variation not captured by other environmental covariates (Arce Guillen et al., 2023). This unexplained spatial variation could reflect an unmeasured driver of movement (e.g., predation risk, conspecific interactions) or centres of attraction (e.g., kill sites, dens), which if unaccounted for, may bias selection parameters for other covariates. In this paper, we will use a very general definition of random effects that includes penalised smooths, which have great flexibility in capturing spatial, group-level, and temporal variation in movement and selection patterns (Hodges, 2014; Wood, 2017).

It is most common to implement SSFs using software for fitting conditional logistic regression models (Signer et al., 2019; Therneau, 2022). Typically, this approach has been limited to relatively simple log-linear models (although see `survival` for non-linear p-spline implementation; Therneau, 2022). Many of the extensions mentioned above have required the use of custom code or specialised packages, such as `inlabru` for spatial random effects (Arce Guillen et al., 2023) and `glmmTMB` for random slope models (Muff et al., 2020). Therefore, the increasing diversity of methodological extensions and implementation methods can present a challenge for practitioners, and it may not be easy to explore model formulations in the same inferential framework.

In this paper, we explain how an SSF can be formulated as a “general smooth model” (i.e., a generalised additive model (GAM) without an exponential family distribution; Wood et al., 2016), and can conveniently be implemented in the popular R package `mgcv`. This framework greatly increases the flexibility of current SSF models, and we will focus on the following: i) modelling non-linear patterns of selection and non-parametric movement kernels, ii) capturing inter-group variability (i.e., random slopes and hierarchical smooths), iii) accounting for unexplained spatial variation via spatial random effects, and iv) assessing selection patterns that change through time and/or space using varying-coefficient models. We will show that this implementation provides a unifying framework to fit, compare, and contrast complex SSF formulations with non-linear and random effects.

## 2 General model formulation and implementation

### 2.1 Step selection functions (SSFs)

Consider a track of two-dimensional animal locations {***s***_1_, ***s***_2_, …, ***s***_*T*_ } collected at regular time intervals. In an SSF, the likelihood of a step ending at location ***s***_*t*+1_ given the previous locations ***s***_1:*t*_ = {***s***_1_, ***s***_2_, … ***s***_*t*_} is

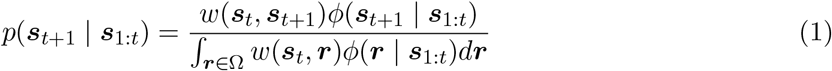

where *w* measures habitat selection, *ϕ* describes the movement patterns of the animal, and Ω is the study area (Rhodes et al., 2005; Forester et al., 2009). The denominator is a normalising constant, such that Equation 1 is a probability density function with respect to the endpoint of the step, ***s***_*t*+1_. The product of *w* and *ϕ* is usually called the SSF, and it is most common to define both components as log-linear models. Then the habitat selection is defined as 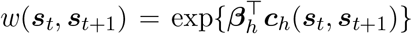 where ***c***_*h*_(***s***_*t*_, ***s***_*t*+1_) is a vector of habitat covariates with associated coefficients ***β***_*h*_. Likewise, the movement kernel is commonly written as 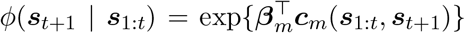 with movement covariates ***c***_*m*_(***s***_1:*t*_, ***s***_*t*+1_) and coefficients ***β***_*m*_. The two exponential functions can be factorised, such that Equation 1 becomes

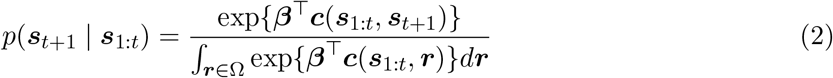

where ***c***(***s***_1:*t*_, ***s***_*t*+1_) = (***c***_*h*_(***s***_*t*_, ***s***_*t*+1_), ***c***_*m*_(***s***_1:*t*_, ***s***_*t*+1_)) and ***β*** = (***β***_*h*_, ***β***_*m*_). Typically ***c***_*h*_(***s***_*t*_, ***s***_*t*+1_) will include spatial covariates (e.g., foraging resources, proxies of risk, etc.), and the parameters ***β***_*h*_ measure the animal’s selection for (positive coefficients) or avoidance of (negative coefficients) these environmental features. A common movement model is the correlated random walk (CRW), and it can be implemented within the SSF framework by including specific functions of the step length and turning angle as covariates in ***c***_*m*_ (Forester et al., 2009; Duchesne et al., 2015; Avgar et al., 2016). The coefficients ***β***_*m*_ are then related to the parameters of the estimated distributions of the movement variables. This approach can be used to model turning angles with a von Mises distribution (Duchesne et al., 2015), and there are several options for step lengths, including the exponential, gamma, log-normal, and Weibull distributions (Forester et al., 2009; Avgar et al., 2016). SSFs can also include interaction terms (i.e., covariate-dependent coefficients) to capture spatiotemporal variation (Forester et al., 2009; Richter et al., 2020; Avgar et al., 2016). Note that some SSFs do not estimate movement jointly with habitat selection (e.g., Fortin et al., 2005), but we focus on SSFs where both *ϕ* and *w* are estimated (sometimes termed an “integrated” SSF; Forester et al., 2009; Avgar et al., 2016).

The normalisation constant of the SSF (i.e., the integral in the denominator) is analytically intractable, and a common approach is to approximate the integral as the sum of function evaluations at random points from some distribution *h* (Michelot et al., 2023). This approximation is nearly likelihood-equivalent to conditional logisitic regression (CLR), and it can be convenient to re-write the SSF as such. That is, in this approach, we restrict the animal’s movement choices to the random and observed points, and we can write the model in terms of the covariate values at these discrete choices. We define a stratum as the observation and random points at each time *t*. Then ***X***_*t*_ is the design matrix, with columns for evaluations of ***c*** (i.e., habitat and movement variables) and one row for each location in the *t*-th stratum (i.e., the observation and the *N* random locations). Assuming the number of random points *N* is constant over all strata, we can approximate the SSF by,

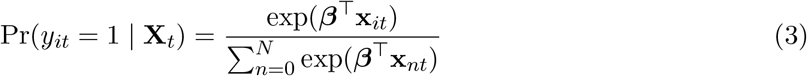

where **x**_*it*_ is the row of **X**_*t*_ indexed by *i*, and

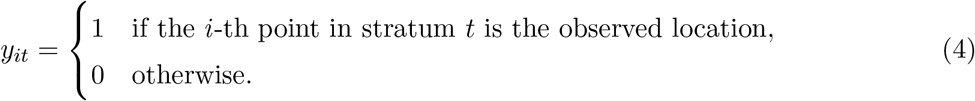

Note that in Equation 3, we temporarily ignored the distribution of random points *h*, and we must account for our sampling via a post hoc correction or “update” to the parameters (Forester et al., 2009; Avgar et al., 2016). Typically random points will be sampled from step length and turning angle distributions, derived from the empirical data (see Section 3.1 and Table 1 for details on sampling and corrections). This approach allows the SSF to be fitted using existing software for CLR (Forester et al., 2009; Avgar et al., 2016), and there are several options for log-linear SSFs, including the R packages `survival` (Therneau, 2022) and `glmmTMB` (Muff et al., 2020). Here, we recognise that Equation 3 is also likelihood-equivalent to a special case of a Cox proportional hazards (Cox PH) model (see Appendix A for details), and can therefore be fitted in the `mgcv` package. This is the same equivalence utilised in `survival`, where CLR can be fitted with `clogit` as a wrapper to `cox.ph` function. Implementation in `mgcv` as a GAM-like general smooth model allows for more flexible formulations with both non-linear and random effects, with model complexity controlled by a data-driven penalty. We describe this modelling framework next.

**Table 1:**
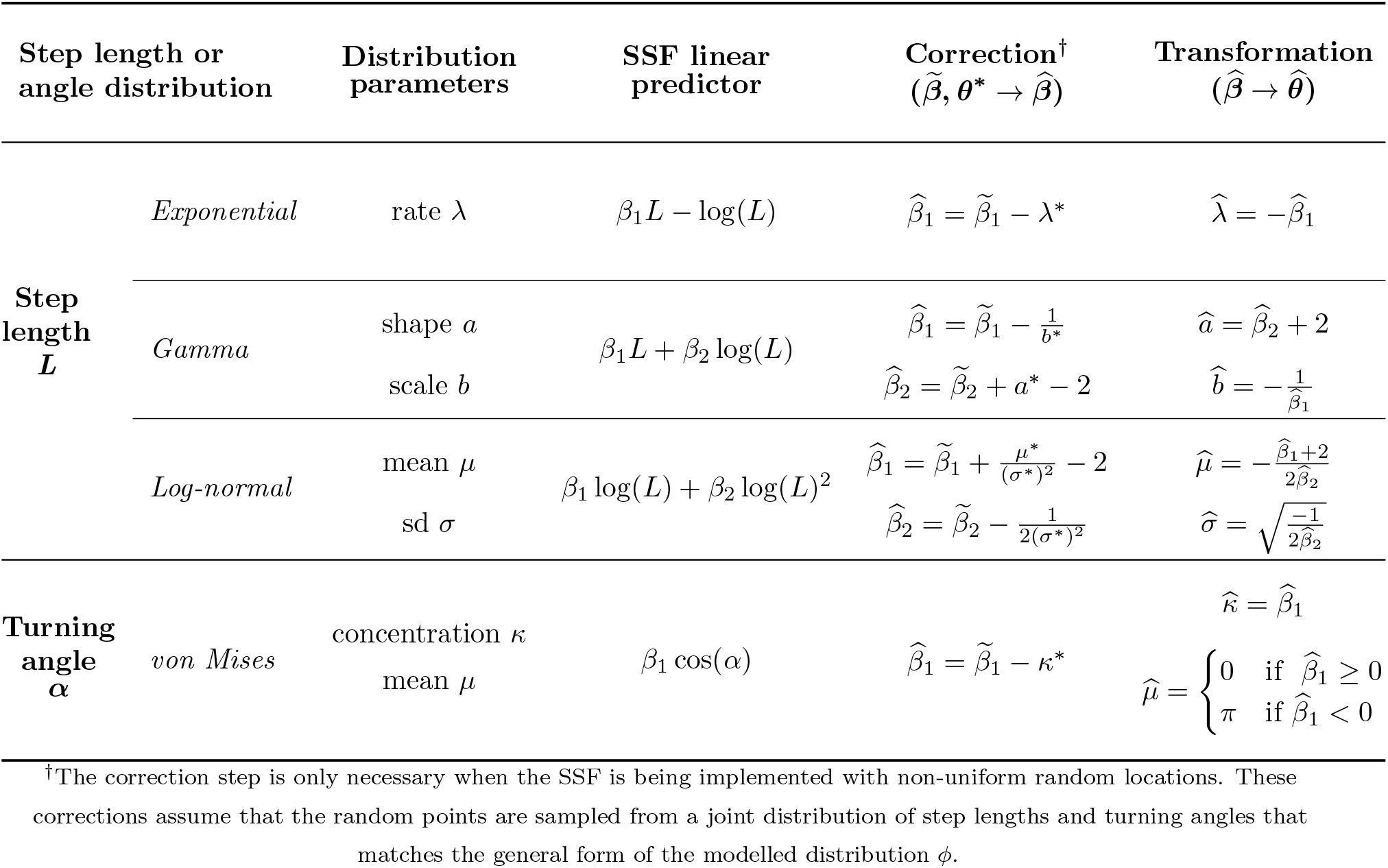
Common step length (L) and turning angle (α) distributions used in SSFs. If the random points are generated from tentative distributions of step lengths and turning angles, we obtain tentative estimates of the coefficients (denoted by 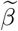). These estimates do not correspond to the parameters of the SSF linear predictor, and therefore, we must correct these parameters (i.e., account for the random point distribution with parameters **θ**^*^) to obtain the SSF parameters (denoted as 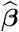), which can then transformed to the distribution parameters of interest 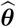.

### 2.2 SSFs with smooth effects

Equation 3 can be extended to include non-linear and “conventional” random effects (e.g., random slopes), which we will collectively call “smooth” terms following the terminology used for GAMs (Wood, 2017). We write the general model as

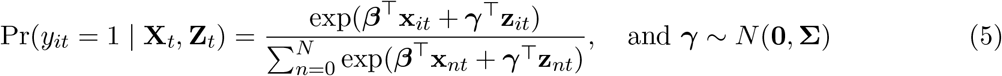

where **X**_*t*_ is the design matrix of fixed effect (unpenalised) terms and **Z**_*t*_ is the design matrix of smooth terms for the stratum *t* (with *i*-th row ***z***_*it*_). Note that this model does not include an intercept, as this would not be identifiable given Equation 5. In this mixed effect formulation, ***β*** is the vector of fixed effects and ***γ*** is the vector of random effects, assumed to follow a multivariate normal distribution with covariance matrix **Σ**. The choice of a normally distributed ***γ*** is sometimes described as a prior, reflecting our belief about the random effect variance or function smoothness (which unperpins a Bayesian view of smoothing and uncertainty quantification; Kimeldorf and Wahba, 1970; Miller, 2021). If **Z**_*t*_ represents basis functions from a single smooth term, then the covariance matrix in Equation 5 is given by **Σ** = **S**^−^*/λ*, where **S** is called the penalty matrix of the smoother (with pseudo-inverse **S**^−^) and *λ* is a smoothing parameter. If **Z**_*t*_ includes more than one smoother or smoothers are penalised by multiple terms, then **Σ**^***−***^ will be a block-diagonal matrix, with the penalty matrices multiplied by penalty-specific smoothing parameters on the matrix diagonal (Wood, 2017).

Equation 5 is very flexible, as mixed effect models provide a convenient framework to describe not only inter-group heterogeneity (e.g., using random slopes), but also non-linear effects using penalised splines and spatial dependence via random fields (Wood, 2017; Hodges, 2014). These formulations correspond to different choices of the covariates in the model matrix **Z**_*t*_ and of the penalty matrix **S** (and therefore the covariance matrix **Σ**). We describe random slopes and random intercepts as “conventional” random effects, but note that throughout the paper we use the term random effects more generally to also describe other smooth effects. Next, we will formalise this equivalence between conventional random effects and penalised smooths, and explain their uses within the SSF framework (see Figure 1 for a summary of key examples). Throughout, we denote a linear predictor for the *i*-th location of stratum *t* as *η*_*it*_ = ***β***^⊤^**x**_*it*_ + ***γ***^⊤^**z**_*it*_ (as in Equation 5).

**Figure 1:**
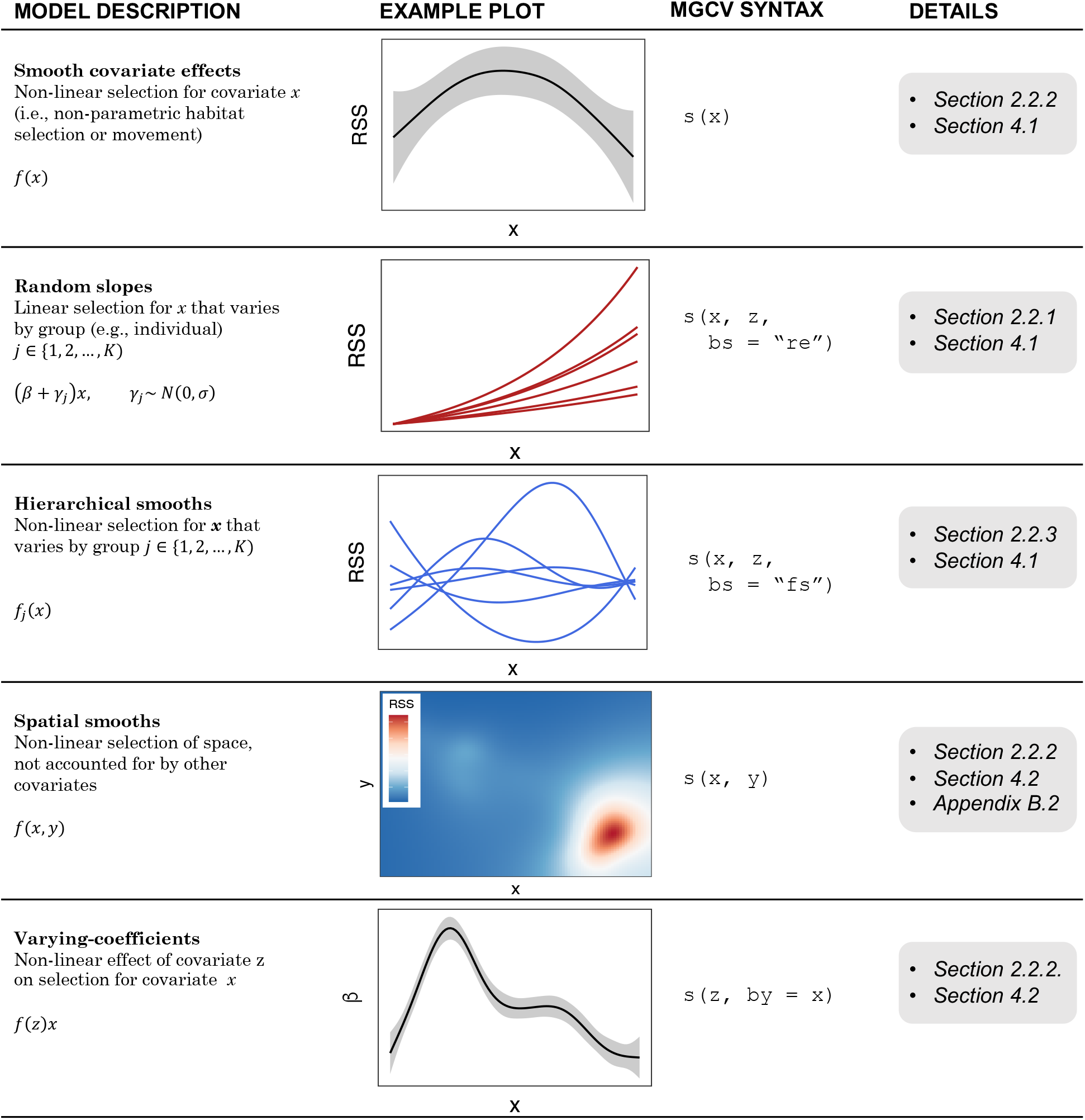
Summary of smooth effects discussed in this paper. “Model description” explains the form of the smooth effect, which is demonstrated visually in the example plot. The mgcv syntax only refers to the relevant term in the model formula, where s() denotes a smooth function, bs is the basis function type, and by is used to specify interactions. Further details of each effect can be found elsewhere in the paper and appendices.

#### 2.2.1 Conventional random effects

“Conventional” random effects, which are used to capture differences between individuals or groups, are perhaps the simplest special case of the mixed effect model in Equation 5. SSFs lack an intercept, and so we will focus on defining a model with random slopes to capture variability in habitat selection or movement patterns (Muff et al., 2020; Chatterjee et al., 2023). The linear predictor of an SSF with independent random slopes for a covariate *x* takes the form,

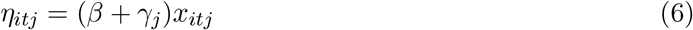

with random effect levels *j* ∈ 1, 2, …, *K*. In this case, each *z*_*it*_ in Equation 5 is zero when the observation is not part of the *j*-th random effect level (and *z*_*it*_ = *x*_*it*_ when it is). The penalty matrix of the i.i.d. random effects is the identity matrix (**S** = **I**), leading to the random effect distribution 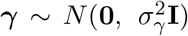, where 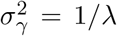. That is, the smoothing parameter *λ* is inversely related to the variance of the random effects; a large value of *λ* corresponds to a small value of 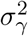 and thus, shrinkage of individual or group-specific parameters toward the population mean (see Appendix B.1 for a simulated example and a short comparison to `glmmTMB`).

#### 2.2.2 Smooth effects

It is possible to model non-linear terms (e.g., penalised splines, Gaussian processes) as random effects. We define a single-penalty smooth function of some covariate *u* as

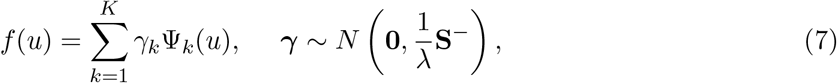

where the Ψ_*k*_ are simple mathematical functions called “basis functions”, weighted by the coefficients *γ*_*k*_ (see Figure 2 for an example; Hefley et al., 2017; Wood, 2017; Pedersen et al., 2019). In Equation 7, the penalty matrix is determined by the choice of smoothing basis, and the penalties encode the assumptions about the prior probability of different function shapes (for e.g, the default thin plate regression splines in `mgcv` assigns lower prior values to functions that have a large squared second derivative; Wood, 2017). The smoothness of the function (i.e., the degree of penalisation) is controlled by *λ*. When *λ* is high, this indicates low covariance of the basis coefficients, such that adjacent basis function evaluations are more similar, resulting in a smoother function (Figure 2). In contrast to simpler polynomials and regression splines with complexity specified *a priori*, we estimate *λ* from the data to control the trade-off between model fit and complexity.

**Figure 2:**
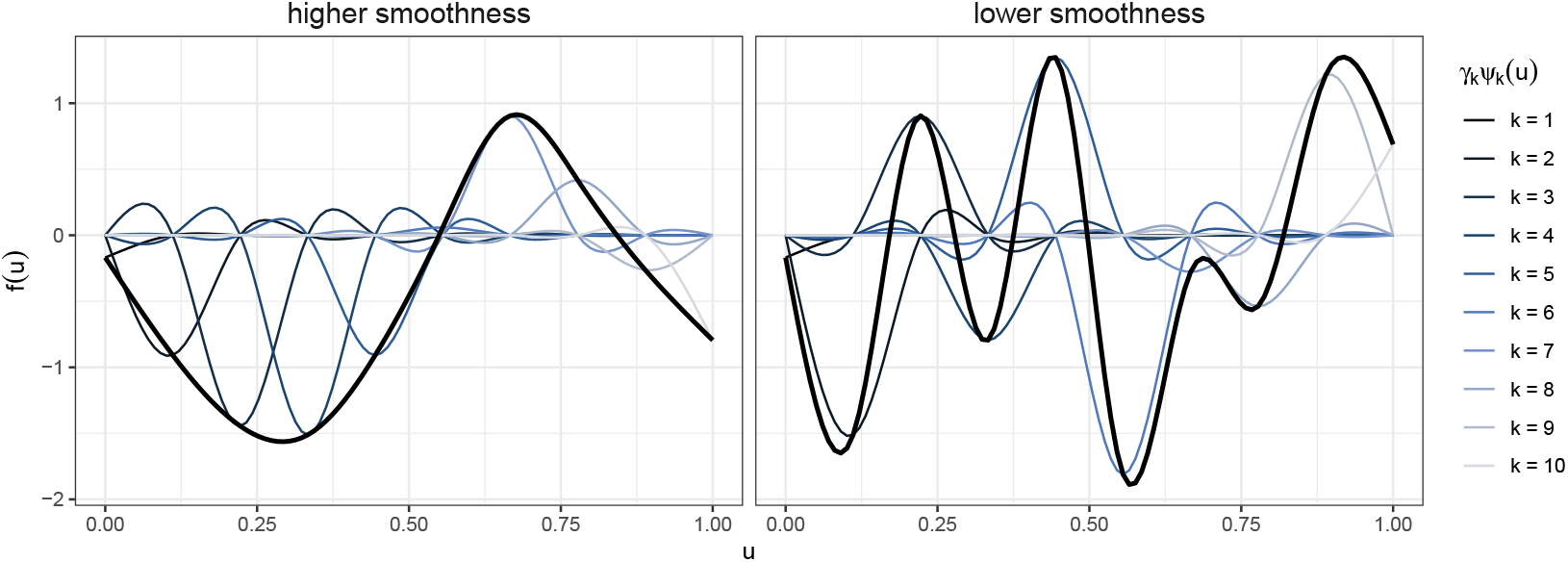
Example of how to derive cubic regression splines: the blue lines are the basis functions **Ψ** multiplied by their coefficients **γ**, which are then summed to obtain the smooth functions (black lines). The panels show examples with different smoothing penalties (i.e., λ).

This approach is also extensible to smooths with *m* ≥ 2 penalties (e.g., adaptive smoothers with variable smoothness along the covariate range) by replacing **S**^−^*/λ* in Equation 7 with the pseudoinverse of a sum of the *m* penalty matrices for a given smoother multiplied by specific penalty terms: 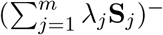. This is important because it allows for modelling non-linear interactions between covariates with different units as smooth terms via tensor-product smoothers (Wood et al., 2013).

There are several applications of smoothing splines, including modelling non-linear covariate effects, random fields, and formulating interaction models with smooth effects. Here, we briefly summarise three important uses of smooth functions in SSFs (which we explore in more detail in Sections 3 and 4):

1. Including smooth functions of covariates can capture non-linear patterns of habitat selection and non-parametric movement kernels. The linear predictor of an SSF with a smooth of covariate *x* (e.g., step length, turning angle, or a spatial feature) takes the form *η*_*it*_ = *f* (*x*_*it*_). This can relax the assumption that animals have a constant rate of selection (see Section 4.1) and simple unimodal distributions of movement variables.
2. Incorporating a spatial smooth (i.e., random effect or random field) can account for spatial variation that is not explained by the other environmental covariates in the model (Arce Guillen et al., 2023). An SSF with a spatial smooth of the spatial coordinates (here, denoted *u* and *v*) has the linear predictor *η*_*it*_ = *f* (*u*_*it*_, *v*_*it*_) with smooth function 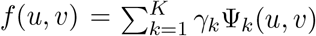, where Ψ_*k*_ is a two-dimensional basis function (Wood, 2017; Hefley et al., 2017). In Appendix B.2, we show how an unknown centre of attraction can be captured with a spatial smooth, and in Section 4.2, we use a spatial smooth to model patterns of a zebra’s space use that cannot be explained by preference for a measured habitat variable.
3. Formulating a model with varying coefficients (i.e., where the linear effect *β* of a covariate varies smoothly with another covariate; Wood, 2017) greatly increases the flexibility of SSFs. An interaction between the covariate *x* and a non-linear effect of *u* can be written as *η*_*it*_ = *f* (*u*_*it*_)*x*_*it*_, where *f* (*u*) is given by Equation 7. That is, the selection coefficient for *x* is specified as a non-linear function of *u*. This approach is highly flexible, as it allows for any specification of non-linear covariate-dependent habitat selection or movement. For example, in Section 4.2, we show how zebra movement speed varies cyclically with time of day, reflecting underlying behavioural variation (Klappstein et al., 2023).

#### 2.2.3 Hierarchical smooths

We can formulate step selection models with multiple levels of smooth effects via hierarchical smooths (Pedersen et al., 2019). In this framework, a separate smooth relationship is estimated for each random effect level (e.g., each individual). This is conceptually similar to a random slope model, as it allows individuals or groups to differ in their selection or movement patterns, but in a non-linear manner. Consider a model with a hierarchical smooth with *K* groups for a continuous covariate *x*. The linear predictor for the *j*-th group is *η*_*itj*_ = *f*_*j*_(*x*_*itj*_), where each smooth *f*_*j*_ takes the general form presented in Equation 7. The same general formulation could be extended to account for inter-group variability in any smooth effect (e.g., spatial smooths). Further, *f*_*j*_ is often modelled as the sum of a population-level smooth relationship and individual-specific smooth deviations from the population. This gives great flexibility to specify which model components are shared across individuals and which are not (e.g., shape or smoothness of the relationship), and Pedersen et al. (2019) describe several important possible formulations. The `sz` smoother has also recently been added to `mgcv` as a type of hierarchical smoother with the main effect factored out of the smooth term, leaving only differences between individuals in the smoother. This allows for estimation of both an overall main effect (via a non-hierarchical smoother) and individual-level effects (via an `sz` smoother). This addresses some of the issues identified in Pedersen et al. (2019) with estimating hierarchical GAM models that include both a global and group-level smoother for the same term.

## 3 Practical guidance

In this section, we provide some practical guidance for practitioners interested in using `mgcv` to fit SSFs with smooth effects. We cover how to sample random points for different movement models, how to choose appropriate settings in `mgcv`, as well as model selection, interpretation, and diagnostics (Figure 3 presents a summary of the workflow).

**Figure 3:**
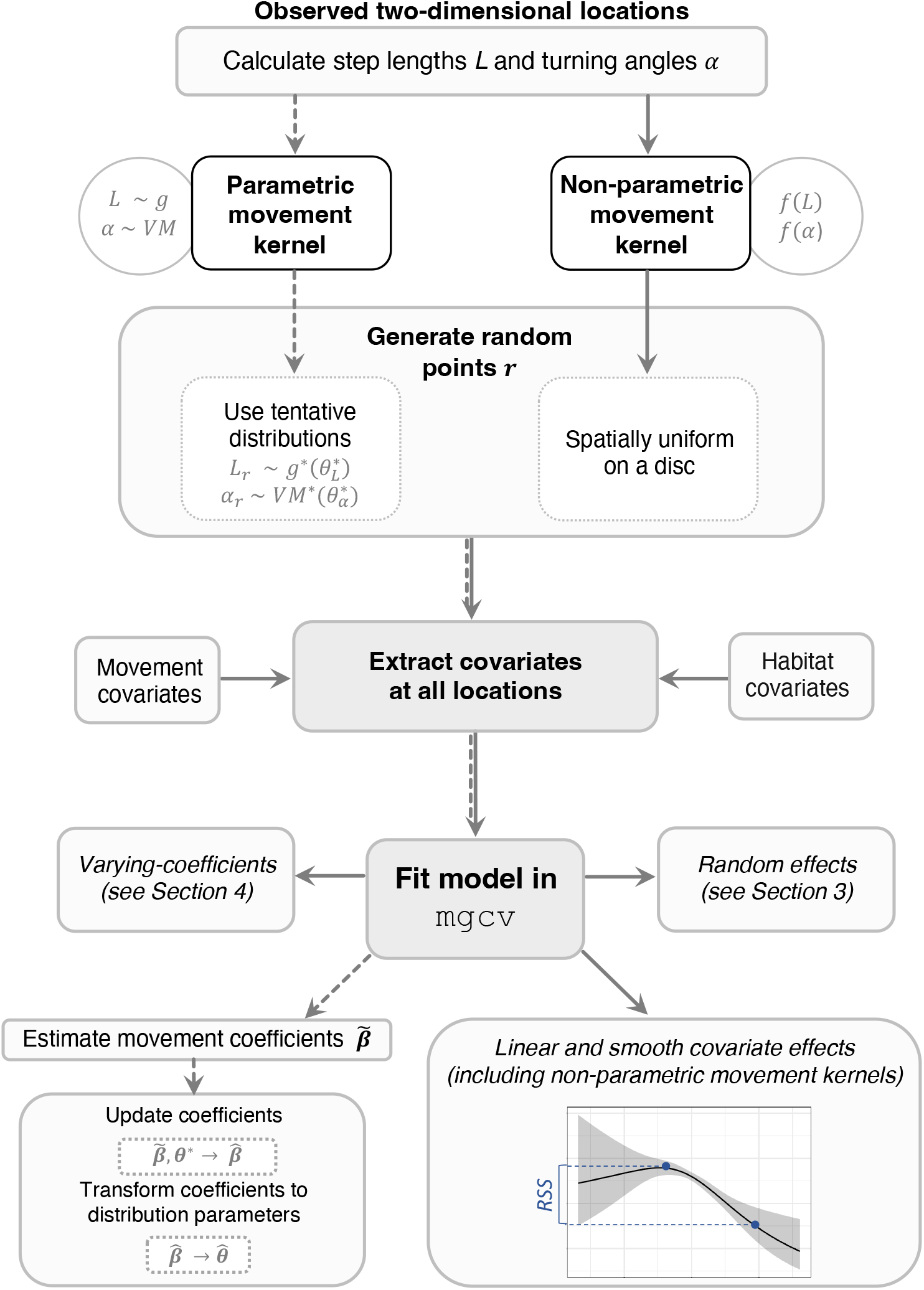
Summary of workflow to implement SSFs described in this paper (dashed lines indicate the most common workflow for parametric movement kernels). L denotes step length, α denotes turning angles, and the distributions g^*^ (any step length distribution from the exponential family) and V M ^*^ (von Mises) used to generate random points are typically estimated from empirical data. RSS is the relative selection strength calculated as the ratio between the two blue points, where the y-axis is the exponential of the partial effect.

### 3.1 Sampling random points

As explained in Section 2.1, the implementation of SSFs requires sampling random points from a two-dimensional distribution *h* (typically based on step lengths and turning angles) to obtain an approximation of the likelihood (Michelot et al., 2023). The choice of *h* is constrained within the CLR framework, and it must usually either have the same general form as *ϕ* or be spatially uniform. For parametric movement kernels, there are post hoc corrections to account for *h* (i.e., to “update” the tentative parameter estimates; Avgar et al., 2016, Table 1), but there is no easy way to apply the same workflow to model non-parametric movement kernels. To estimate a non-parametric movement kernel, we therefore recommend sampling points {***r***_*t*1_, …, ***r***_*tN*_ } uniformly over a disc centred on the previous observed location ***s***_*t*−1_, with a radius *R* close to the maximum observed step length for adequate spatial coverage (Klappstein et al., 2022; Michelot et al., 2023). In practice, this can be done by generating each turning angle from Unif(−*π, π*) and each step length as the square root of a random draw from Unif(0, *R*^2^).

Although spatially uniform points are flexible, they are generally not the most efficient sampling scheme. For parametric movement kernels, a better method is to simulate more random points in areas where the animal is more likely to move (i.e., importance sampling; Avgar et al., 2016; Michelot et al., 2023). For this purpose, *h* is often chosen as the spatial distribution implied by some step length and turning angle distributions. As we are implementing the SSF as a Cox PH model, we must then correct (or “update”) the estimated parameters to obtain estimates of the actual movement parameters of *ϕ* (see Table 1 for common step and turning angle distributions with the relevant covariates and corrections; Avgar et al., 2016). Note that if points are sampled uniformly in space, *h* is constant and does not need to be corrected for.

### 3.2 Implementation in mgcv

Using `mgcv` requires implementing an SSF as a Cox PH model via the `cox.ph` family. To do so, we define a constant “event time” variable (`times`; all observations set to the same value) in conjunction with a stratification variable (`stratum`; an identifier for each grouping of an observed location and its random points) as our response, and use the `weights` argument to distinguish the observed location from the random points (i.e., via an indicator variable; `obs`). Ultimately, this defines a CLR model, and the linear predictor can be defined using standard formula syntax in `mgcv`. For example, a model with non-parametric distributions of step lengths and turning angles, and a linear effect of habitat covariate *x* is specified as,

~~~
fit <-gam (cbind (times, stratum) ∼ s(step) + s(angle) + x,
 data = dataset,
 family = cox. ph,
 weights = obs)
~~~

where `times` is a vector of 1’s, and `obs` is a vector of 0’s and 1’s. The model parameters are estimated via Restricted Maximum Likelhood (REML).

#### 3.2.1 Choice of basis functions and basis dimension

The choice of a smoother defines the form of the basis functions and the penalty term that controls the function smoothness (Wood, 2017; Pedersen et al., 2019). Therefore, various random effect and smooth models can be fitted simply by choosing the appropriate basis functions. By default, `mgcv` uses thin plate regression splines for both one- and multi-dimensional smooths. Although these generally perform well, there are several other general smoother options available (see Wood, 2017, and `mgcv` documentation for full descriptions). For example, a spatial random field could be specified as a Gaussian process with several options for the covariance function. As outlined by Miller et al. (2020), this approach is similar to the stochastic partial differential equation approach, which has been previously implemented for spatial random effects in SSFs via `inlabru` (Arce Guillen et al., 2023). Specialised smoothers can be used to better account for boundary behaviour (e.g., soap film smoothers, Duchon splines) and non-isotropic coordinates (e.g., splines on the sphere). Additionally, cyclic cubic regression splines can be used to capture cyclic patterns, which are ubiquitous in ecology (e.g., effects of time of day; Feldmann et al., 2023). Since we can view a random effect as a smooth (and vice versa), we can also specify a random slope model via the basis functions (see Figure 1 and coded examples on GitHub for relevant settings in `mgcv`).

In principle, the basis dimension *K* (i.e., number of columns in the associated design matrix) only sets the upper limit to the wiggliness of the function. That is, for large enough *K*, the smoothness of the function is constrained by the penalty matrices and estimated smoothness parameters *λ*. In practice, the shape of the estimated function might therefore not be affected much by this choice, but a large *K* will be more computationally demanding (Wood, 2017). We suggest trying multiple choices of *K* to assess if model outputs change between fits, and checking how *K* compares to the effective degrees of freedom (if the values are similar, it may indicate that a higher *K* is required). Note that in the case of random slopes, *K* is the number of groups and cannot be changed.

### 3.3 Model interpretation

Both smooth and parametric (main) effects can be interpreted in terms of relative selection strength (RSS; Avgar et al., 2017; Fieberg et al., 2021). For linear effects, there is an intuitive interpretation of the coefficient, where (holding all else equal) exp(*β*) is how many times more likely an animal is to take a step when the covariate increases by 1 (similarly to other regression coefficients). The relative selection strength between any two steps can be calculated as the ratio of the predicted selection exp(*βx*_*i*_) for each step (and can be calculated in the `amt` package; Signer et al., 2019). For example, consider steps with covariate values *x*_1_ and *x*_2_; the RSS of the step with *x*_1_ is *RSS*(*x*_1_, *x*_2_) = exp(*βx*_1_)*/* exp(*βx*_2_). For a smooth term, we lose the straightforward interpretation of exp(*β*), but we can calculate the RSS in the same way. It may also be useful to plot the predicted selection for a grid of covariate values, but note that the *y*-axis can only be interpreted relatively (Figures 3, 4, 5). Note that this same interpretation applies to spatial smooths, but where the RSS is calculated between two spatial locations (i.e., the spatial smooth should not be confused with the spatial distribution of the animal; Figure 4b). Note that interaction terms (e.g., varying coefficients) and random effects have standard interpretations (as described in Wood, 2017)

**Figure 4:**
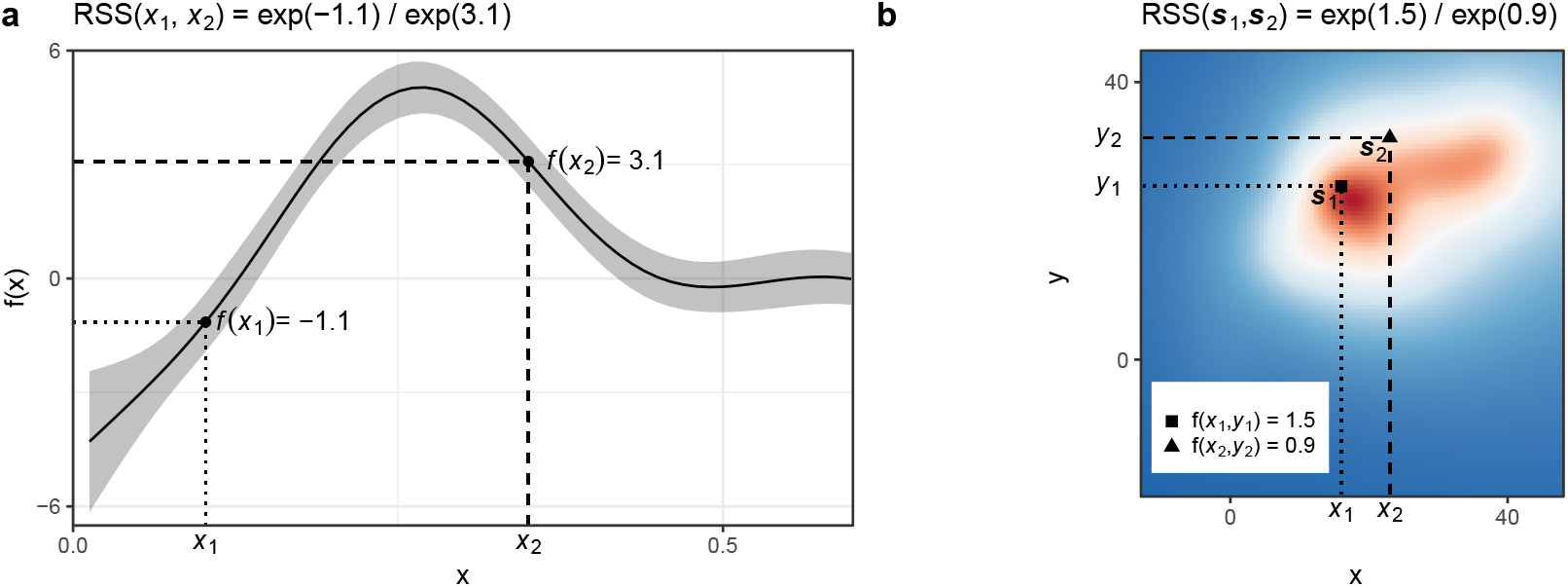
Relative selection strength (RSS) examples. (a) shows the RSS for the covariate x (all else held equal), where RSS(x_1_, x_2_) refers to the RSS of location with covariate value x_1_ relative to x_2_. (b) shows how to interpret a spatial smooth, where the RSS is based on spatial locations (i.e., different values of easting x and northing y), where RSS(**s**_1_, **s**_2_) refers to the RSS of the first location **s**_1_ (with coordinates x_1_, y_1_) compared to the second **s**_2_ (with coordinates x_2_, y_2_).

**Figure 5:**
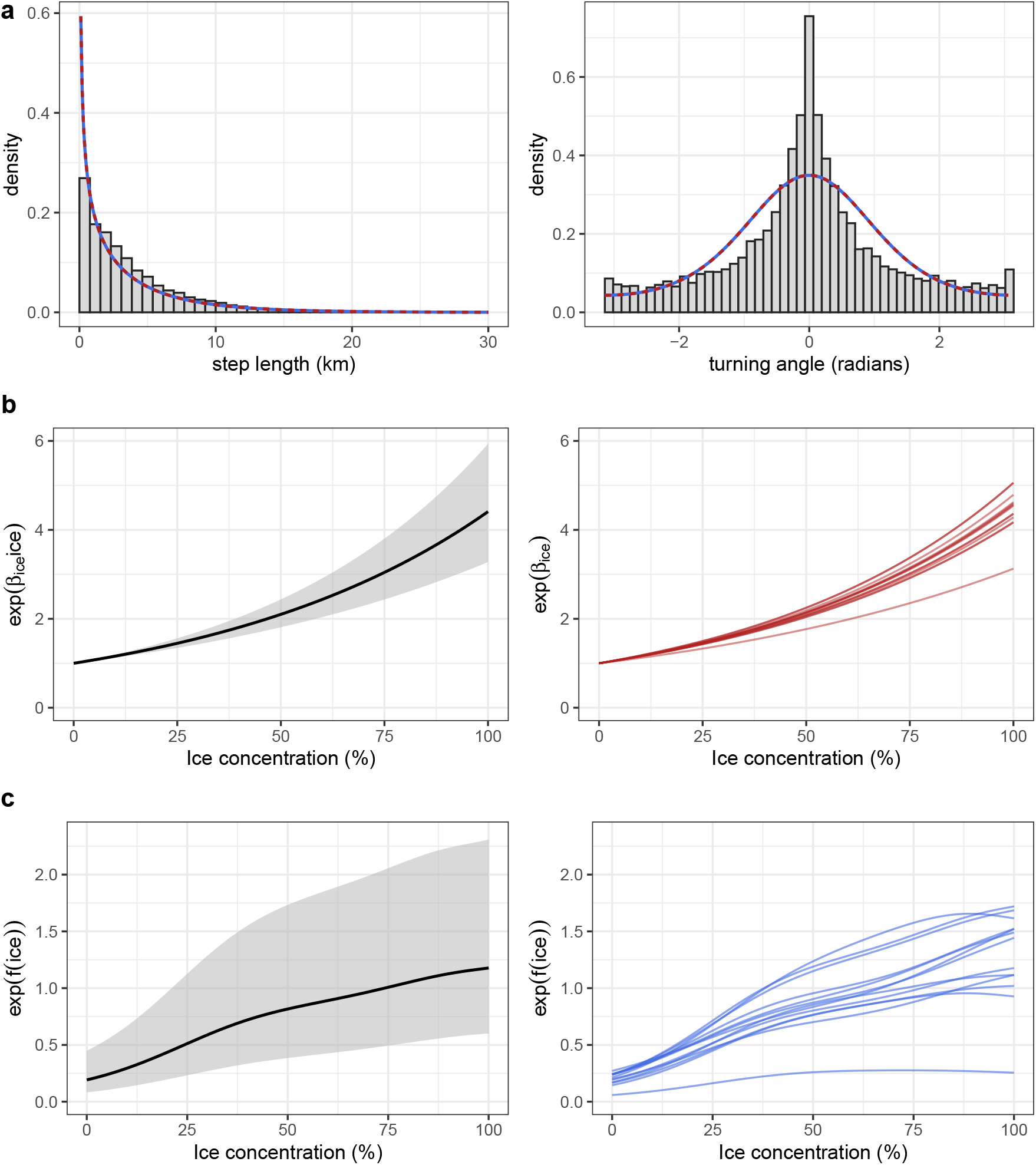
Results for polar bear example: (a) step length and turning angle distributions for both models (shown in dashed/solid lines), (b) random slopes for ice concentration, and (c) hierarchical smooths for ice concentration. In both (b) and (c), the left plot shows the population estimate with 95% CIs and right plot shows individual-level estimates.

Confidence intervals for estimated smooth curves should be interpreted as pointwise (rather than simultaneous) CIs for a given smooth curve (Wood, 2017). Confidence intervals for smooth curves estimated using REML have been shown to have good coverage properties when averaged across the function, but might have above- or below-nominal coverage at different points in the curve, so should be interpreted with caution. Visualising posterior simulations from the fitted curve provides a better visual guide to the degree of functional uncertainty in a fitted model, rather than just the estimated curve and confidence interval. These posterior simulations can be produced via the `smooth samples` function of the `gratia` package (Simpson, 2023).

### 3.4 Model selection and diagnostics

AIC is commonly used for variable selection in regression models. There are two types of AIC for models with random effects fitted with `mgcv`: i) marginal AIC (which tends to favour complex models) and ii) conditional AIC (favouring simpler models) (Wood, 2017). Wood et al. (2016) describes a “corrected” conditional AIC, which better accounts for smoothing parameter uncertainty in the penalty term. This is the default when `AIC()` is applied to a `gam` model object, and it is one possible approach to choose among competing model formulations.

Another approach is to use penalisation to automatically exclude smooth terms that have no clear effect, by “shrinking” the effect to zero. For most smooths, increasing *λ* to a large number only constrains the relationship to be any straight line, rather than zero (the set of straight lines is called the null space of the penalty). This problem can be resolved by also penalising non-zero straight lines. In `mgcv`, this can be done with specialised shrinkage bases, for which the smoothing parameter *λ* also penalises the null space (i.e., where *λ* → ∞ effectively removes the corresponding term from the model), or by including an additional penalty with the `select=TRUE` option for any smooth term (i.e., the “double-penalty” approach; Marra and Wood, 2011).

In general, there are no simple residual-based checks for SSFs, and this is not solved by our proposed implementation in `mgcv`. Although `mgcv` has a convenient `gam.check` function, the residuals are specifically designed to assess the assumptions of the Cox PH model. For example, Schoen-field residuals test the proportional hazards assumption and deviance residuals are derived from Martingale residuals based on the cumulative hazard function (Wood, 2017). However, neither of these are assumptions of an SSF, and therefore we advise against using these residuals for model checking. Diagnositics are notoriously difficult for SSFs, and although out of the scope of this paper, simulation-based methods could be promising for SSFs with smooths (Fieberg et al., 2018; DiRenzo et al., 2023; Fieberg et al., 2023).

## 4 Illustrative examples

In this section, we present two real data examples to illustrate the models and implementation described in Section 2: i) we explore how to best account for inter-individual variability in polar bear (*Ursus maritimus*) habitat selection, and ii) we formulate a model with a spatial smooth and a time-varying (parametric) movement kernel for a single plains zebra (*Equus quagga*). For both examples, we matched each observed location with 25 random steps, generated from a gamma distribution of step lengths (with parameters derived from the empirical data) and uniform turning angles. In all models, we modelled step lengths *L* with a gamma distribution (by including *L* and log(*L*) as covariates) and turning angles *α* with a von Mises distribution (by including cos(*α*) as a covariate).

### 4.1 Capturing inter-individual variability of polar bear habitat selection

We obtained 4-hour GPS locations from 13 polar bears in the Beaufort Sea. We regularised the movement tracks and interpolated spatial covariates (following Klappstein et al., 2020, 2022). We analysed a total of 14,927 locations (range: 862 − 1810 per individual). The main goal of inference was to assess inter-individual variability in selection for ice concentration, which is linked to polar bear prey distribution and energetic costs of travel (Pilfold et al., 2014; Klappstein et al., 2022). Therefore, we considered two models that allow for this inter-group variation: a model with random slopes and a model with hierarchical smooths. We assumed that the movement kernel *ϕ* (described at the beginning of Section 4) was shared across all individuals. The linear predictor for the *i*-th location of stratum *t* of the random slope model (for *j* ∈ 1, 2, … 13 individuals) takes the form,

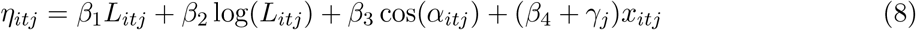

where *x* is the ice concentration. The analogous hierarchical smooth model has the form,

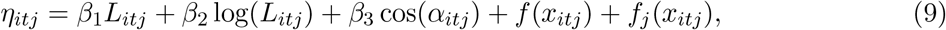

where both smooth functions had *K* = 5 basis functions. Each model contained both a global term for the slope or smooth, as well as inter-individual deviations from that global mean, and the main difference was that selection for ice concentration was either modelled as a log-linear effect (Equation 8), or as a smooth (i.e., non-linear) effect (Equation 9).

Estimates of movement parameters were the same (to two decimal places) for both models 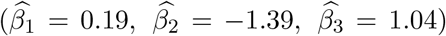, indicating that polar bear step lengths followed a gamma distribution with mean 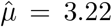 and standard deviation 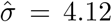, and turning angles followed a von Mises distribution with mean 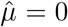 and concentration 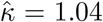 (Figure 5a). Note that the animals’ movement patterns are affected by habitat selection, so the observed and estimated distributions in Figure 5a are not necessarily expected to match completely. The models differed in their characterisation of the selection for ice concentration. The population mean of the random slopes model indicated that the “typical” bear selected for higher values of ice concentration, and was 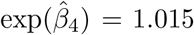 times more likely to take a step for every 1% increase in ice concentration. We cannot easily derive this relationship from the hierarchical smooth model, but we can assess relative selection between any two competing ice concentrations by comparing the estimated smooth function at each point. For example, consider if the average bear was presented with a choice of 0% or 50% ice concentration. The smooth and linear models predict that the bear would be 4.3 and 2.1 (respectively) times more likely to choose the step with 50%. However, note that the smooth function varies across the covariate range and the RSS will not be the same for any two points with 50% difference (as is the case for a linear model).

Both models captured inter-individual variability, but the smooths model also captured variability in the pattern of selection (i.e., individuals have functions with different shapes; Figure 5c). We compared the models with AIC (as described in Wood et al., 2016) and found that the hierarchical smooth model had the lower AIC (ΔAIC = 38.3). This is consistent with previous results on polar bear habitat selection, which indicated that individual polar bears select for an optimal ice concentration between 60 to 100% (rather than a log-linear relationship; Klappstein et al., 2022) corresponding with maximum prey biomass at 85% (Pilfold et al., 2014).

### 4.2 Spatiotemporal variation of zebra movement and habitat selection

We analysed 7246 GPS locations from a single plains zebra, collected at a 30-minute resolution in Hwange National Park in Zimbabwe from January–April 2014 (previously described in Michelot et al., 2020; Klappstein et al., 2023). The goal of the analysis was threefold: i) assess habitat selection for different vegetation types, ii) capture temporal variation in movement patterns, and iii) account for any remaining spatial variation using a spatial smooth.

We assessed habitat selection for a categorical vegetation variable (woodland, bushland, bushed grassland, and grassland as the reference category). As in the previous example, we defined *ϕ* with a gamma step length distribution and a von Mises distribution of turning angles. We allowed the scale parameter of the gamma distribution (i.e., the step length coefficient) to vary based on the time of day *τ* via a cyclic spline (with *K* = 10), to capture temporal patterns in movement speed that repeat or occur each day. Lastly, we included a spatial smooth (i.e., an isometric smooth interaction between the easting *u* and northing *v*) using two-dimensional thin plate regression splines (with default *K* = 30). The model linear predictor was,

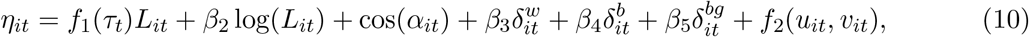

where *δ*^*w*^, *δ*^*b*^, *δ*^*bg*^ are indicator variables for woodland, bushland, and bushed grassland.

The zebra’s mean step length changed throughout the day (captured by variability in the associated coefficient; Figure 6a), ranging from approximately 80-100m in the evening to 200-250m in the early morning and mid-afternoon (Figure 6b). Consistent with previous findings, our results indicate that the zebra selected for grassland over all other habitat types (Figure 6c; Michelot et al., 2020; Klappstein et al., 2023). The spatial smooth showed remaining spatial pattern, once other covariates had been considered. In particular, there was one area at the northern edge of the map that the zebra disproportionately selected for (Figure 6d), which could indicate an unaccounted for spatial feature that the zebra was attracted to. Note that the estimated spatial smooth should be interpreted in terms of RSS (see Figure 4b) and cannot be interpreted as a spatial distribution.

**Figure 6:**
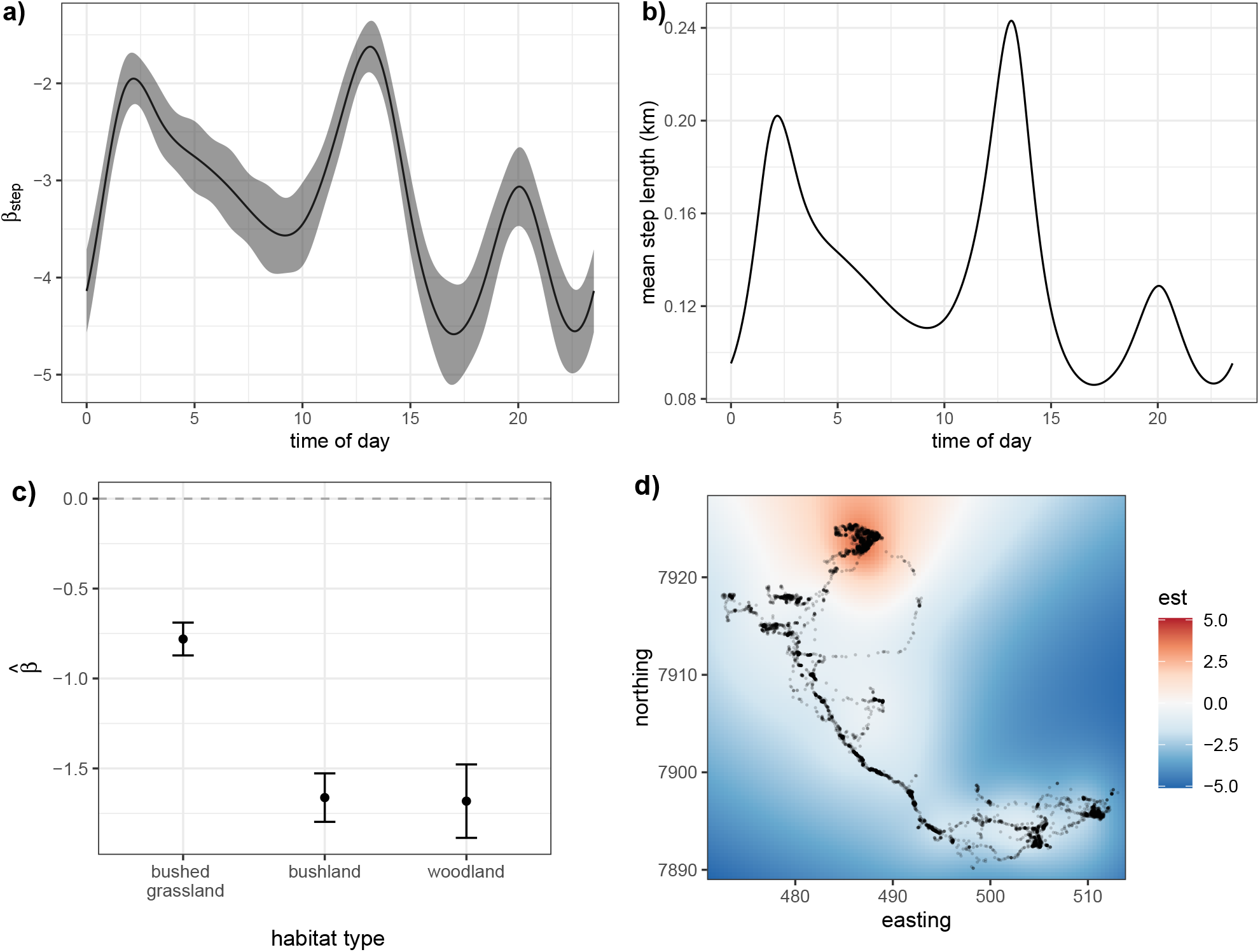
Results of the zebra model. (a) and (b) show how step length varies as a function of time of day. (a) shows how the the step length coefficient varied throughout the day, and (b) is the same relationship, translated to the scale of the mean step length. (c) Selection coefficients with 95% CIs for habitat types (with grassland as a reference category, i.e., corresponding to zero). (d) Partial effect of the spatial smooth (on the log scale).

## 5 Discussion

In this paper, we explained how smooth (i.e., random) effects provide a unifying framework for many SSF extensions. Using smooths, we can account for non-linearity, inter-individual variability, and other spatial and temporal dependencies in habitat selection and movement patterns. This approach can be easily implemented in the widely-used R package `mgcv`, and we gave a general overview of the flexible nature of smooth effects. We focused on several key cases we believe will be particularly relevant, but there are many additional possibilities that could further extend the capabilities of SSA. We hope this framework encourages practitioners to think about their SSF formulations more flexibly, via the inclusion of interactions, random effects, and more realistic movement dynamics.

In general, smooth interactions are powerful tools in SSA, as they can model complex covariate-dependence in habitat selection and movement. We showed how varying-coefficient models can capture daily patterns of zebra movement speed. This approach can be generalised to other SSF covariates, where any time-varying coefficient could capture phases of movement or habitat selection corresponding to behavioural changes. This accomplishes a similar goal as state-switching SSFs (Nicosia et al., 2017; Klappstein et al., 2023), but rather than discrete behavioural switches, movement or selection patterns are allowed to vary smoothly and continuously through time (similar to approaches in Hanks et al., 2015; Michelot et al., 2021). We could extend this approach to model selection coefficients that vary over space if there is reason to believe that animals may modulate their behaviour in response to environmental features (e.g., in response to conspecifics; Smith et al., 2023). These time- or space-varying coefficients could be accomplished with specific covariates (e.g., time of day, season, habitat covariates), or with general temporal or spatial smooth interactions (e.g., a spatially varying coefficient model; Comber et al., 2022). Although we focused on “varying coefficients”, which refers to allowing a *linear* effect to vary smoothly with another covariate, smooth effects can also interact smoothly with other covariates. Interactions between covariates on different scales are possible with tensor products, which can include the marginal effects of one or both variables (Wood, 2017). For example, this could capture a non-parametric movement kernel with dependence between step lengths and turning angles (similar to copula-based kernels; Hodel and Fieberg, 2022).

It is often of interest to capture inter-group (e.g., inter-individual) variability in movement or habitat selection patterns. For this purpose, random slopes have been adopted for SSFs, but these are limited to linear habitat selection and parametric movement (Duchesne et al., 2010; Muff et al., 2020). We showed how simple random slopes can be fitted as a smooth term, and how we can improve on this linear framework by incorporating hierarchical smooths into the SSF framework. Hierarchical smooths are more flexible and can capture inter-group variability in *non-linear* (i.e., smooth) terms. McCabe et al. (2021) explored the utility of hierarchical smooths in the context of large-scale species distributions across multiple individuals via RSFs, which do not explicitly model movement. By extending hierarchical smooths to SSFs, we can further investigate inter-individual differences in both animal habitat selection *and* movement patterns that vary through time and space (Chatterjee et al., 2023). This framework also makes it straightforward to account for inter-individual variability via including individual-specific covariates, via continuous or “factor-smooth” interactions (e.g., animals may change their habitat selection throughout their life cycle or there may be inter-sex differences).

Although smooth effects afford modellers a vast range of options, this additional flexibility comes with more computational and estimation challenges. Model fitting may be less numerically stable and more sensitive to the choice of random points (i.e., integration methods; Michelot et al., 2023). Depending on the basis dimension, there can be many parameters to estimate, adding computational burden, and requiring trial and error to appropriately specify *K*. Model checking is still difficult, and future work could explore simulation-based checks which have recently been proposed for simpler SSFs (Fieberg et al., 2018, 2023). In general, confounding may become a larger issue as more general random effects and interactions are incorporated into an SSF. For example, although spatial smoothing is a powerful way to account for unexplained spatial variation, there is potential for spatial confounding if environmental features are spatially correlated (Hodges and Reich, 2010). In general, practitioners should carefully consider where additional complexity/flexibility is needed in their model, and inspect model outputs for signs of estimation problems (e.g., very large standard errors, convergence issues).

The smooths framework provides a simple way to formulate and fit several recent, complex extensions of step selection analysis. Non-linear effects can be specified without needing to specify model complexity *a priori* (e.g., using polynomials) and more flexible cyclic patterns can be captured without the use of trigonometic functions (Feldmann et al., 2023). Further, random slopes can be implemented with `mgcv`, similar to other approaches in the packages `glmmTMB` or `inlabru` (Muff et al., 2020). Arce Guillen et al. (2023) also used `inlabru` to implement spatial random effects for SSFs. In `mgcv`, it is possible to specify spatial smooths with several forms, including Gaussian processes (to match previous approaches; Arce Guillen et al., 2023; Miller et al., 2020). Our proposed implementation allows individual-specific and spatial random effects to be combined with other smooth terms described in this paper, using software for GAMs that many practitioners are already familiar with (Wood, 2017; Pedersen et al., 2019).

## Data availability

Code and data are available at GitHub (https://github.com/NJKlappstein/smoothSSF).

## Acknowledgements

We thank Andrew Derocher for providing the polar bear data and Simon ChamaillÉ-Jammes for providing the zebra data used in the illustrative examples. Also, we thank Tal Avgar for taking the time to discuss the project in its early stages.

## A Relationship to a Cox proportional hazards model

A Cox proportional hazards model is typically used for time-to-event data (often the event is the subject’s death). The hazards ratio for subject *i* with event at time *t* is

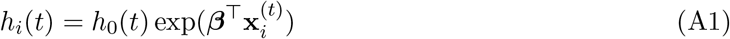

where *h*_0_(*t*) is the baseline hazard, which can be factored out. The partial likelihood of the model for *T* subjects is,

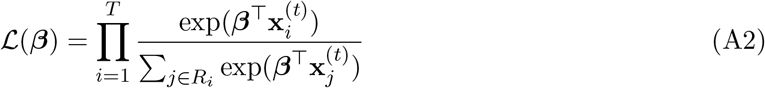

where *R*_*i*_ is the risk set consisting of all subjects that are alive at time *t* (including the subject). Importantly, *R*_*i*_ also contains any censored subjects (i.e., subjects whose last recorded time was not their death/event), but these do not have their own hazard ratio included (i.e., they are not included in the subject list *i* ∈ 1, 2, …, *T*). The Cox model can also be stratified, in which each risk set is only considered over a particular grouping (e.g., only subjects from the same hospital).

Conditional logistic regression (and by extension, SSA) can be fitted as a special case of a Cox model with a particular data structure (Therneau, 2022). In this case, each time step with a matched set of an observed location and control locations are considered both the stratum and the subject. The control locations are set as the censored set (i.e., their SSF is only used to calculate the likelihood of the case in that stratum). Then we can write the Cox PH likelihood equivalent to an SSF (i.e., CLR; with a few changes to the notation),

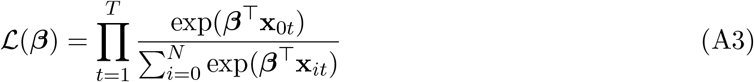

where *x*_0*t*_ are the covariates of the observed location at time *t, N* is the number of random points, and *T* is the number of observed locations in the dataset.

### Extension to smooth effects

The full (un-penalised) track likelihood of an SSF with smooth effects is,

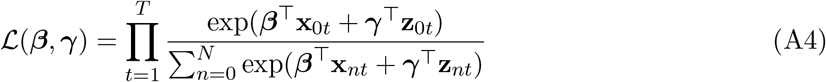

where *T* is the total number of observations (i.e., stratum), and **x**_0*t*_ and **z**_0*t*_ are the rows of the design matrices corresponding to the case (i.e., observed) locations. To implement a *penalised* smooth, we can add a penalty term to the likelihood in Equation A4 based on the smooth coefficients and penalty matrix. The degree of the penalisation is determined by a smoothing parameter *λ*, such that the full penalised log-likelihood is,

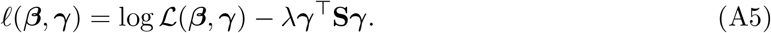

## B Simulations

This section provides several simulations to assess the performance and capabilities of smoothing within the context of SSFs in `mgcv`. All simulations followed the same algorithm to produce a movement track (also described in Klappstein et al., 2022; Michelot et al., 2023):

1. Generate starting location ***s***_1_ randomly from within the middle 50% of the study area Ω
2. To generate each ***s***_*t*_ in {***s***_2_, ***s***_3_, …, ***s***_*T*_ }
  a. Generate many possible endpoints {***r***_1_, ***r***_2_, …, ***r***_*J*_ } uniformly on a disc centred on ***s***_*t*_ (with a pre-defined radius), and evaluate/interpolate relevant covariates.
  b. Select ***s***_*t*+1_ from the possible endpoints with probabilities given by the SSF

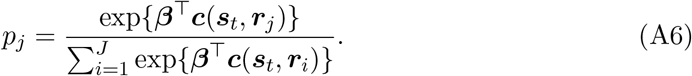

For all simulations, we used *J* = 1000 possible endpoints on a disc with radii given as the 99th percentile of the step length distribution. We focus on three main simulations: i) comparing `mgcv` to `glmmTMB` for estimating random slopes, ii) showing how spatial smoothing can be used to account for an “unknown” centre of attraction, and iii) demonstrating how a non-parametric movement kernel and non-linear habitat effects can be used to capture dynamics that arise from and underlying state-switching process.

### B.1 Random slope models in mgcv **and** glmmTMB

Muff et al. (2020) proposed a method to include random slopes in an SSF using a conditional Poisson model with a large fixed variance to estimate stratum-specific intercepts in the `R` packages `glmmTMB`. Our approach accomplishes the same goal, but because the model accounts for stratification through the CLR form, it is unnecessary to estimate stratum intercepts. However, the function `gam()` will be less efficient at estimating many random effect levels than `glmmTMB` as it is unable to exploit the highly sparse nature of the random effect penalty matrix, and there is no option to use other `mgcv` options (e.g., `gamm` or `bam`) with the `cox.ph` family. Here, we compare how well `glmmTMB` and `mgcv` estimate (known) SSF parameters with random slopes, and compare fitting times. Note that this only represents a simple simulation scenario and is not meant to be an exhaustive or conclusive comparison between packages.

#### Simulating and fitting data

Using the simulation algorithm presented above, we generated 100 movement tracks of length *T* = 2500. Step lengths *L* followed a gamma distribution with shape *a* = 2 and rate *b* = −*β*_*step*_ = 4. We randomly generated a covariate (denoted *cov*) with spatial autocorrelation *ρ* = 2 (for details, see Michelot et al., 2023). We simulated selection for the habitat covariate with random slopes. We simulated the *i*-th individual’s random slope as *β*_*i,cov*_ ∼ *N* (*β*_*cov*_, *σ*), where the population-level selection was set to *β*_*cov*_ = 5 with standard deviation *σ* = 0.5. We fitted each track with `mgcv` using the model term `s(cov, ID, bs = “re”)` for random slopes, and `glmmTMB` (following Muff et al., 2020).

#### Simulation results

Figure B1 shows all estimates. The estimates of fixed effects were very similar between `mgcv` and `glmmTMB`: the coefficient for the habitat covariate (*β*_*cov*_) was consistently overestimated and the step length coefficient (*β*_*step*_) was generally estimated well. However, the packages differed in the estimates for the random effect variance. `glmmTMB` tended to have wider variability across the estimates (which increased with the number of individuals), with some negative bias (consistent with Muff et al., 2020). The median variance estimates from `mgcv` were closer to the truth and there was less variability between estimates. Fitting was much faster in `glmmTMB`, particular for many individuals.

**Figure B1:**
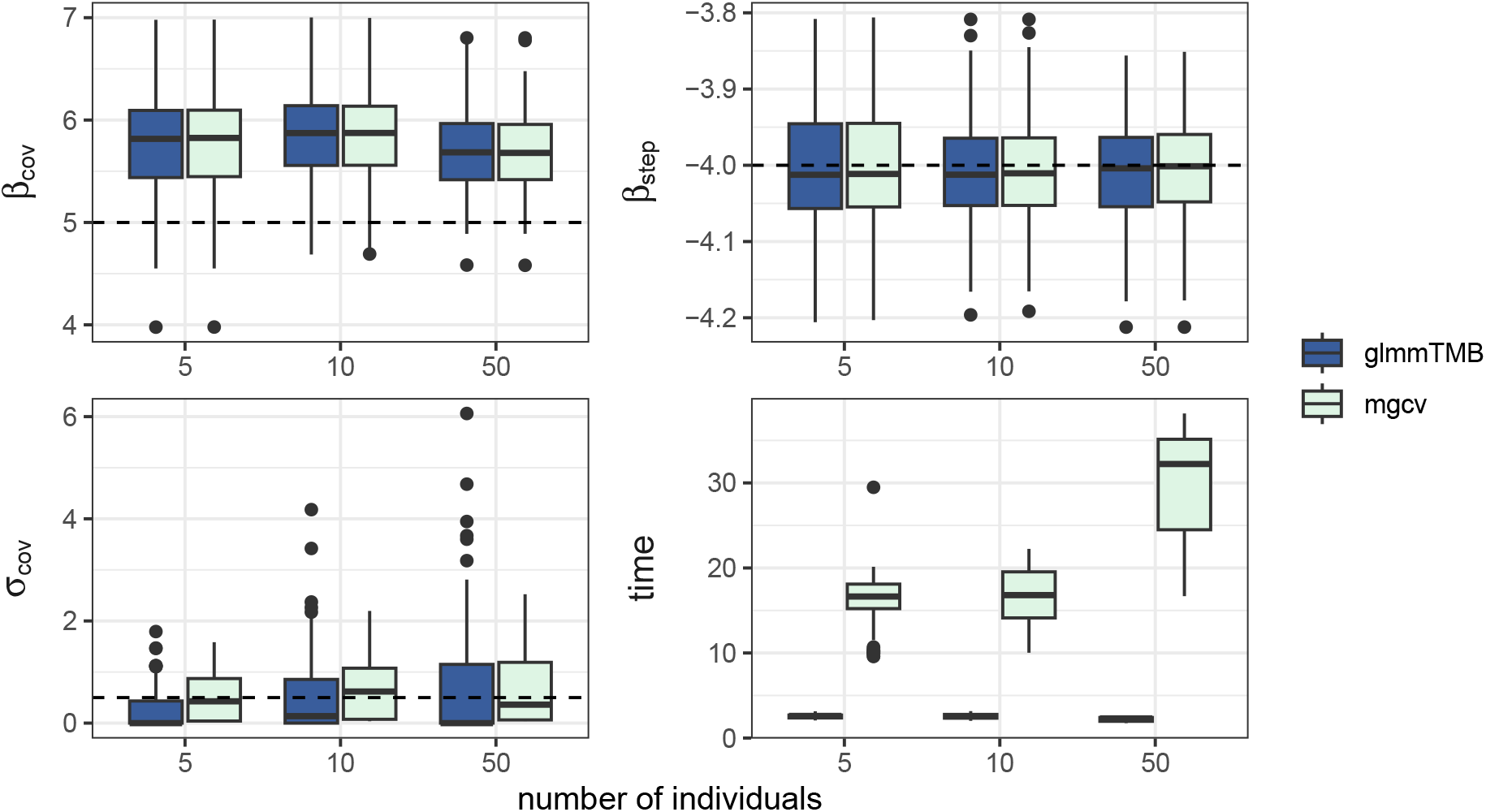
Results from simulation study, where `glmmTMB` and mgcv were used to fit SSFs with random slopes to 300 movement tracks (100 each had 5, 10, and 50 individuals) of length T = 2500. β_step_ is the SSF coefficient for step length; β_cov_ is the population-level selection coefficient for the habitat covariate and σ_cov_ is the standard deviation of the random slope distribution; and time is the fitting time in seconds. In the first three panels, the dashed line is the true parameter value.

### B.2 Spatial smoothing

Here, we show how spatial smoothing can be used to capture an unaccounted for centre for attraction. We simulated a movement track of length *T* = 2000 with selection for a habitat covariate (*β*_*cov*_ = 5) and distance to a centre of attraction (*β*_*c*_ = −0.075). Step lengths *L* followed a gamma distribution with shape *a* = 2 and rate *b* = −*β*_*step*_ = 1.6. When we fitted the model, we included the habitat covariate, but omitted the centre of attraction. Instead, we included a general spatial smooth to capture the effect of the centre of attraction. The spatial smooth is able to accurately capture the relative preference for movement toward the centre of attraction, with no *a priori* information on where that centre is.

**Figure B2:**
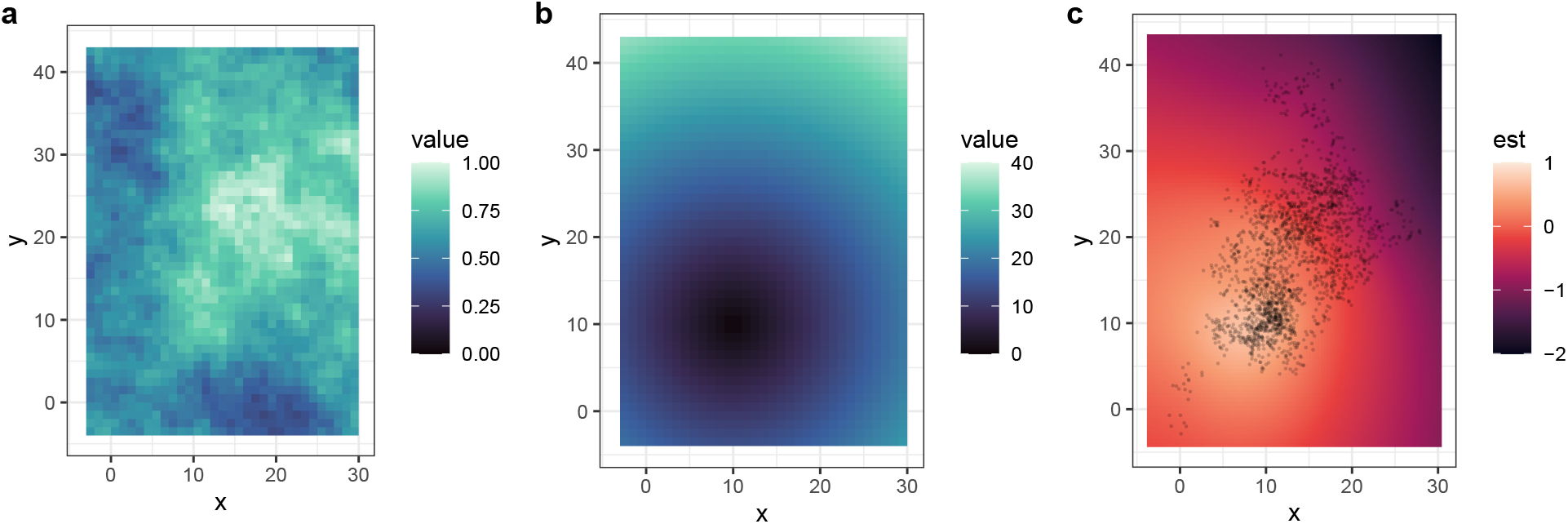
Location data were simulated from an SSF with selection for a spatial covariate (a) and distance to a centre of attraction (b). The right panel (c) shows the estimated spatial smooth when covariate (a) was included, but the distance covariate (b) was omitted. Dots are the simulated locations.

## References

Arce Guillen, R., Lindgren, F., Muff, S., Glass, T. W., Breed, G. A., and Schlägel, U. E. (2023). Accounting for unobserved spatial variation in step selection analyses of animal movement via spatial random effects. Methods in Ecology and Evolution, 14(10):2639–2653.

Avgar, T., Lele, S. R., Keim, J. L., and Boyce, M. S. (2017). Relative Selection Strength: Quantifying effect size in habitat- and step-selection inference. Ecology and Evolution, 7(14):5322–5330.

Avgar, T., Potts, J. R., Lewis, M. A., and Boyce, M. S. (2016). Integrated step selection analysis: Bridging the gap between resource selection and animal movement. Methods in Ecology and Evolution, 7:619–630.

Chatterjee, N., Wolfson, D., Kim, D., Velez, J., Freeman, S., Bacheler, N. M., Shertzer, K., Taylor, J. C., and Fieberg, J. (2023). Modeling individual variability in habitat selection and movement using integrated step-selection analyses. bioRxiv.

Comber, A., Harris, P., and Brunsdon, C. (2022). Spatially varying coefficient regression with GAM gaussian process splines: GAM(e)-on. AGILE: GIScience Series, 3:1–6.

Dejeante, R., Valeix, M., and ChamaillÉ-Jammes, S. (2023). Time-varying habitat selection analysis: A model and applications for studying diel, seasonal and post-release changes. bioRxiv.

DiRenzo, G. V., Hanks, E., and Miller, D. A. W. (2023). A practical guide to understanding and validating complex models using data simulations. Methods in Ecology and Evolution, 14(1):203–217.

Duchesne, T., Fortin, D., and Courbin, N. (2010). Mixed conditional logistic regression for habitat selection studies. Journal of Animal Ecology, 79(3):548–555.

Duchesne, T., Fortin, D., and Rivest, L.-P. (2015). Equivalence between step selection functions and biased correlated random walks for statistical inference on animal movement. PLoS ONE, 10(4):e0122947.

Feldmann, C. C., Mews, S., Coculla, A., Stanewsky, R., and Langrock, R. (2023). Flexible modelling of diel and other periodic variation in hidden Markov models. Journal of Statistical Theory and Practice, 17(3):45.

Fieberg, J., Forester, J. D., Street, G. M., Johnson, D. H., ArchMiller, A. A., and Matthiopoulos, J. (2018). Used-habitat calibration plots: A new procedure for validating species distribution, resource selection, and step-selection models. Ecography, 41:737–752.

Fieberg, J., Freeman, S., and Signer, J. (2023). Evaluating goodness-of-fit of animal movement models using lineups. bioRxiv, 10.1101/2023.08.10.552754.

Fieberg, J., Signer, J., Smith, B., and Avgar, T. (2021). A ‘How to’ guide for interpreting parameters in habitat-selection analyses. Journal of Animal Ecology, 90(5):1027–1043.

Forester, J., Kyung Im, H., and Rathouz, P. (2009). Accounting for animal movement in estimation of resource selection functions: Sampling and data analysis. Ecology, 90(12):3554–3565.

Fortin, D., Beyer, H. L., Boyce, M. S., Smith, D. W., Duchesne, T., and Mao, J. S. (2005). Wolves influence elk movements: Behaviour shapes a trophic cascade in Yellowstone National Park. Ecology, 86(5):1320–1330.

Hanks, E. M., Hooten, M. B., and Alldredge, M. W. (2015). Continuous-time discrete-space models for animal movement. The Annals of Applied Statistics, 9(1):145–165.

Hefley, T. J., Broms, K. M., Brost, B. M., Buderman, F. E., Kay, S. L., Scharf, H. R., Tipton, J. R., Williams, P. J., and Hooten, M. B. (2017). The basis function approach for modeling autocorrelation in ecological data. Ecology, 98(3):632–646.

Hodel, F. and Fieberg, J. (2022). Circular-linear copulae for animal movement data. Methods in Ecology and Evolution, 13:1001–1013.

Hodges, J. S. (2014). Richly Parameterized Linear Models: Additive, Time Series, and Spatial Models Using Random Effects. Texts in Statistical Sciences. CRC Press, Boca Raton, FL.

Hodges, J. S. and Reich, B. J. (2010). Adding spatially-correlated errors can mess up the fixed effect you love. The American Statistician, 64(4):325–334.

Kimeldorf, G. S. and Wahba, G. (1970). A correspondence between bayesian estimation on stochastic processes and smoothing by splines. The Annals of Mathematical Statistics, 41(2):495–502.

Klappstein, N. J., Potts, J. R., Michelot, T., Börger, L., Pilfold, N. W., Lewis, M. A., and Derocher, A. E. (2022). Energy-based step selection analysis: Modelling the energetic drivers of animal movement and habitat use. Journal of Animal Ecology, 91:946–957.

Klappstein, N. J., Togunov, R. R., Reimer, J. R., Lunn, N. J., and Derocher, A. E. (2020). Patterns of sea ice drift and polar bear (Ursus maritimus) movement in Hudson Bay. Marine Ecology Progress Series, 641:227–240.

Klappstein, NJ., Thomas, L., and Michelot, T. (2023). Flexible hidden Markov models for behaviour-dependent habitat selection. Movement Ecology, 11:30.

Marra, G. and Wood, S. N. (2011). Practical variable selection for generalized additive models. Computational Statistics and Data Analysis, 55:2372–2387.

McCabe, J. D., Clare, J. D., Miller, T. A., Katzner, T. E., Cooper, J., Somershoe, S., Hanni, D., Kelly, C. A., Sargent, R., Soehren, E. C., Threadgill, C., Maddox, M., Stober, J., Martell, M., Salo, T., Berry, A., Lanzone, M. J., Braham, M. A., and McClure, C. J. W. (2021). Resource selection functions based on hierarchical generalized additive models provide new insights into individual animal variation and species distributions. Ecography, 44(12):1756–1768.

Michelot, T., Blackwell, P. G., ChamaillÉ-Jammes, S., and Matthiopoulos, J. (2020). Inference in MCMC step selection models. Biometrics, 76:438–447.

Michelot, T., Glennie, R., Harris, C., and Thomas, L. (2021). Varying-coefficient stochastic differential equations with applications in ecology. Journal of Agricultural, Biological and Environmental Statistics, 26(3):446–463.

Michelot, T., Klappstein, N. J., Potts, J. R., and Fieberg, J. (2023). Understanding step selection analysis through numerical integration. Methods in Ecology and Evolution, 00:1–12.

Miller, D. L. (2021). Bayesian views of generalized additive modelling. arXiv.

Miller, D. L., Glennie, R., and Seaton, A. E. (2020). Understanding the stochastic partial differential equation approach to smoothing. Journal of Agricultural, Biological and Environmental Statistics, 25(1):1–16.

Muff, S., Signer, J., and Fieberg, J. (2020). Accounting for individual-specific variation in habitat-selection studies: Efficient estimation of mixed-effects models using Bayesian or frequentist computation. Journal of Animal Ecology, 89(1):80–92.

Nicosia, A., Duchesne, T., Rivest, L.-P., and Fortin, D. (2017). A multi-state conditional logistic regression model for the analysis of animal movement. The Annals of Applied Statistics, 11(3):1537–1560.

Pedersen, E. J., Miller, D. L., Simpson, G. L., and Ross, N. (2019). Hierarchical generalized additive models in ecology: An introduction with mgcv. PeerJ, 7:e6876.

Pilfold, N. W., Derocher, A. E., and Richardson, E. S. (2014). Influence of intraspecific competition on the distribution of a wide-ranging, non-territorial carnivore. Global Ecology and Biogeography, 23(4):425–435.

Pohle, J., Signer, J., Eccard, J. A., Dammhahn, M., and Schlägel, U. E. (2023). How to account for behavioural states in step-selection analysis: A model comparison. arXiv.

Potts, J. R. and Börger, L. (2022). How to scale up from animal movement decisions to spatiotemporal patterns: An approach via step selection. Journal of Animal Ecology, 00:1–14.

Rhodes, J. R., Mcalpine, C. A., Lunney, D., and Possingham, H. P. (2005). A spatially explicit habitat selection model incorporating home range behavior. Ecology, 86(5):1199–1205.

Richter, L., Balkenhol, N., Raab, C., Reinecke, H., Meißner, M., Herzog, S., Isselstein, J., and Signer, J. (2020). So close and yet so different: The importance of considering temporal dynamics to understand habitat selection. Basic and Applied Ecology, 43:99–109.

Signer, J., Fieberg, J., and Avgar, T. (2017). Estimating utilization distributions from fitted step-selection functions. Ecosphere, 8(4):e01771.

Signer, J., Fieberg, J., and Avgar, T. (2019). Animal movement tools (amt): R package for managing tracking data and conducting habitat selection analyses. Ecology and Evolution, 9:880–890.

Signer, J., Fieberg, J., Reineking, B., Schlägel, U., Smith, B., Balkenhol, N., and Avgar, T. (2023). Simulating animal space use from fitted integrated Step-Selection Functions (iSSF). bioRxiv, page 10.1101/2023.08.10.552754.

Simpson, G. L. (2023). Gratia: Graceful ggplot-based graphics and other functions for GAMs fitted using mgcv. R package version 0.8.1.38.

Smith, B. J., MacNulty, D. R., Stahler, D. R., Smith, D. W., and Avgar, T. (2023). Density-dependent habitat selection alters drivers of population distribution in northern Yellowstone elk. Ecology Letters, 26(2):245–256.

Therneau, T. (2022). A Package for Survival Analysis in R.

Wood, S. N. (2017). Generalized Additive Models: An Introduction with R. Chapman and Hall/CRC.

Wood, S. N., Pya, N., and Säfken, B. (2016). Smoothing parameter and model selection for general smooth models. Journal of the American Statistical Association, 111(516):1548–1563.

Wood, S. N., Scheipl, F., and Faraway, J. J. (2013). Straightforward intermediate rank tensor product smoothing in mixed models. Statistics and Computing, 23(3):341–360.

